# FlowAtlas.jl: an interactive tool bridging FlowJo with computational tools in Julia

**DOI:** 10.1101/2023.12.21.572741

**Authors:** Valerie Coppard, Grisha Szep, Zoya Georgieva, Sarah K. Howlett, Lorna B. Jarvis, Daniel B. Rainbow, Ondrej Suchanek, Edward J. Needham, Hani S. Mousa, David K. Menon, Felix Feyertag, Krishna T. Mahbubani, Kourosh Saeb-Parsy, Joanne L. Jones

## Abstract

As the dimensionality, throughput, and complexity of cytometry data increases, so does the demand for user-friendly, interactive analysis tools that leverage high-performance machine learning frameworks. Here we introduce FlowAtlas.jl: an interactive web application that bridges the user-friendly environment of FlowJo and computational tools in Julia developed by the scientific machine learning community. We demonstrate the capabilities of FlowAtlas using a novel human multi-tissue, multi-donor immune cell dataset, highlighting key immunological findings.

## Introduction

Rapid advancements in flow and mass cytometry have brought about a new era of high-dimensional cell phenotyping. However, this has not been matched by developments in free-access, coding-free, user-friendly, interactive data analysis tools. Computational pipelines built in scripting languages such as R or Python, require significant coding literacy, hampering their adoption by the wider biomedical community. Additionally, the limited computing power of most laboratory computers often demands data down-sampling to run commonly used dimensionality reduction (DR) methods, risking loss of rare cell populations.

Although DR and cell population clustering algorithms have gradually been integrated into popular analysis platforms such as FCS Express and FlowJo as core features or add-on plugins, these implementations of algorithms such as FlowSOM and tSNE can lack downstream interactivity with the dimensionality-reduced data and still require substantial data down-sampling, ultimately reducing their utility. In addition, as open data access becomes the standard, there is a need for computational tools that allow datasets, acquired using different cytometry panels, to be integrated to facilitate data re-use and validation.

Here we introduce FlowAtlas — a free-access, graphical data analysis environment that aims to address these problems. We chose to write FlowAtlas in Julia [1], a programming language specifically designed for high-performance scientific computing and machine learning applications. This gave us access to some of the fastest algorithms available today [1],[2]. We showcase the capabilities of FlowAtlas using a novel, human flow cytometry dataset, consisting of immune cells extracted from tissues of five deceased organ donors and immunophenotyped using three different antibody panels.

## FlowAtlas Design

### FlowAtlas integrates with FlowJo

We designed FlowAtlas to be an open source, fully graphical, interactive high-dimensional data exploration tool that does not rely on command-line input or coding literacy. FlowAtlas links the familiar FlowJo workflow with a high-performance machine learning framework enabling rapid computation of millions of high-dimensional events without the need for down-sampling (**Figure 1**).

**Figure 1.**
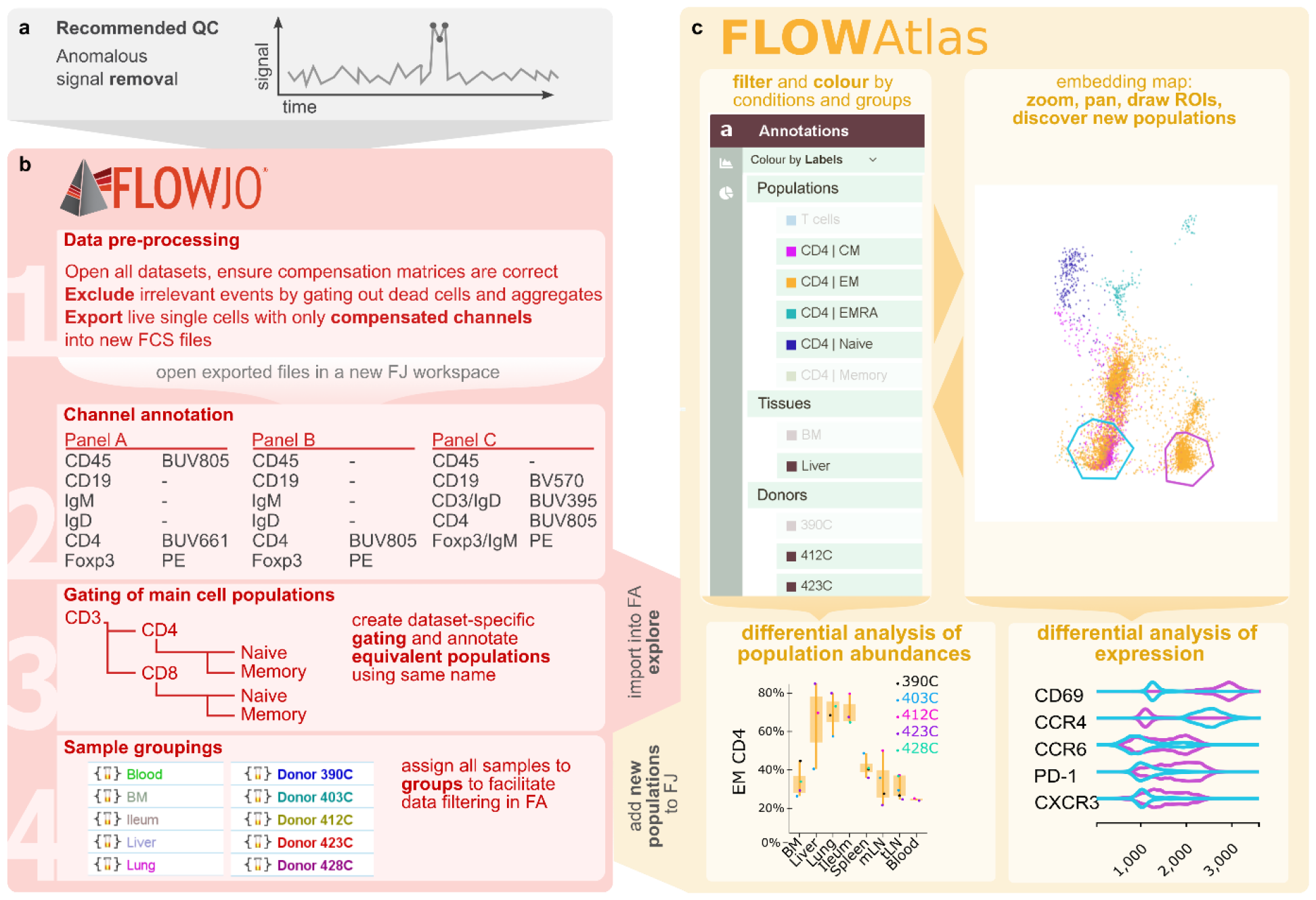
Overview of FlowAtlas workflow with FlowJo. **a**, First, anomalous events are removed from the raw data. **b**, High-quality files are then pre-processed in FlowJo (Step 1) and exported. Exported files are then opened in a new FlowJo workspace and prepared for FlowAtlas-assisted analysis. Step 2: annotation of fluorescence channels; Step 3: panel-specific gating of matched populations across all datasets; Step 4: grouping of samples by desired conditions. Importing the updated workspace file into FlowAtlas triggers automatic panel merging, embedding calculation, and launches interactive web interface (**c**), where embedded events can be re-coloured and filtered by conditions and groups that were defined in FlowJo. ROIs can be drawn directly in the embedding generating violin plots showing marker expression. Box plots can be generated to show frequencies of selected populations and conditions. Novel populations identified in FlowAtlas can be validated and annotated in FlowJo. The updated workspace file can then be reopened in FlowAtlas to import new annotations; FJ - FlowJo; FA - FlowAtlas.

FlowAtlas parses user-defined individual channel transformation settings from FlowJo as well as channel, gate and sample group names, ensuring optimal embedding geometry and ease of data exploration. The resulting embedding is highly interactive, offering zooming to explore deeper cluster structures, colouring and filtering of embedded events by custom conditions, generation of frequency statistics and drawing of regions of interest (ROIs) to perform comparative analysis of marker expression using violin plots. Moreover, FlowAtlas allows merging and concurrent analysis of non-identical panels. Individual samples remain identifiable in the embedding, since the files are not concatenated.

Data exploration happens in an iterative, user-guided discovery loop with FlowJo, where traditional FlowJo gating strategies provide the initial annotation of main cell populations, experimental conditions, and sample groupings to enable the identification of new subpopulations in the interactive embedding. The user periodically returns to FlowJo to add new population annotations as they are discovered in FlowAtlas.

### FlowAtlas enables rapid dimensionality reduction without data downsampling

We eliminated the need for data down-sampling and enabled visual exploration of hundreds of millions of cells by utilising methods within the GigaSOM.jl^3^ library and the interactive web libraries OpenLayers [3] and D3.js [4].

GigaSOM.jl library and its constituent algorithms implement the functionality of self-organising map (SOM)-based clustering [5] and dimensionality reduction in Julia programming language. This enables considerable improvements in computational performance [6] over the current gold-standard SOM and metaclustering-based R package FlowSOM [7], utilised by the majority of open-source analysis workflows and commercial software platforms including FlowJo and Cytobank [8]. The dimensionality reduction algorithm, EmbedSOM, used in the GigaSOM library, has demonstrated a reduction in the computational time requirements of 10-30-fold against popular dimensionality reduction algorithms including UMAP and tSNE [9].

We compared the computational performance of FlowAtlas to two alternative tools for dimensionality reduction that also do not require command-line input on a laboratory laptop with the following identical configuration: Dell XPS15, 64-bit Windows OS, 32GB RAM, 8^th^ generation core i7-8750H processor, 2.20 GHz. Examples graphical outputs from DR with each tool are shown in **Supplementary Figure 1**.

Our tissue-derived immune cell conventional flow cytometry dataset, which is presented as an example throughout this manuscript, consists of 3.88 million total live single cell events (32 FCS files, 19 fluorescence parameters). Donor characteristics, panels and antibodies used are shown in **Supplementary Table 1, Supplementary Table 2** and **Supplementary Table 3**.

Dimensionality reduction of samples stained with panel C (2.32 million events) in FlowJo (v10.8.1), using the inbuilt tSNE function, took 49 min. In FCS Express (v7.18.0025), the same subset of samples was processed in 125 min. The full dataset could not be subjected to DR on these platforms due to panel differences preventing file concatenation.

The full dataset (panel A, B, and C 3.88 million events) was processed in FlowAtlas in 18 min, which included DR and clustering steps (**Table 1**). When analysed as individual files or group of files concatenated by panel, FlowJo tSNE processed the full dataset of 3.88 million events in 6 hours. We did not attempt the same procedure in FCS Express, but it was expected to exceed 125 min required for DR of panel C samples.

**Table 1.**
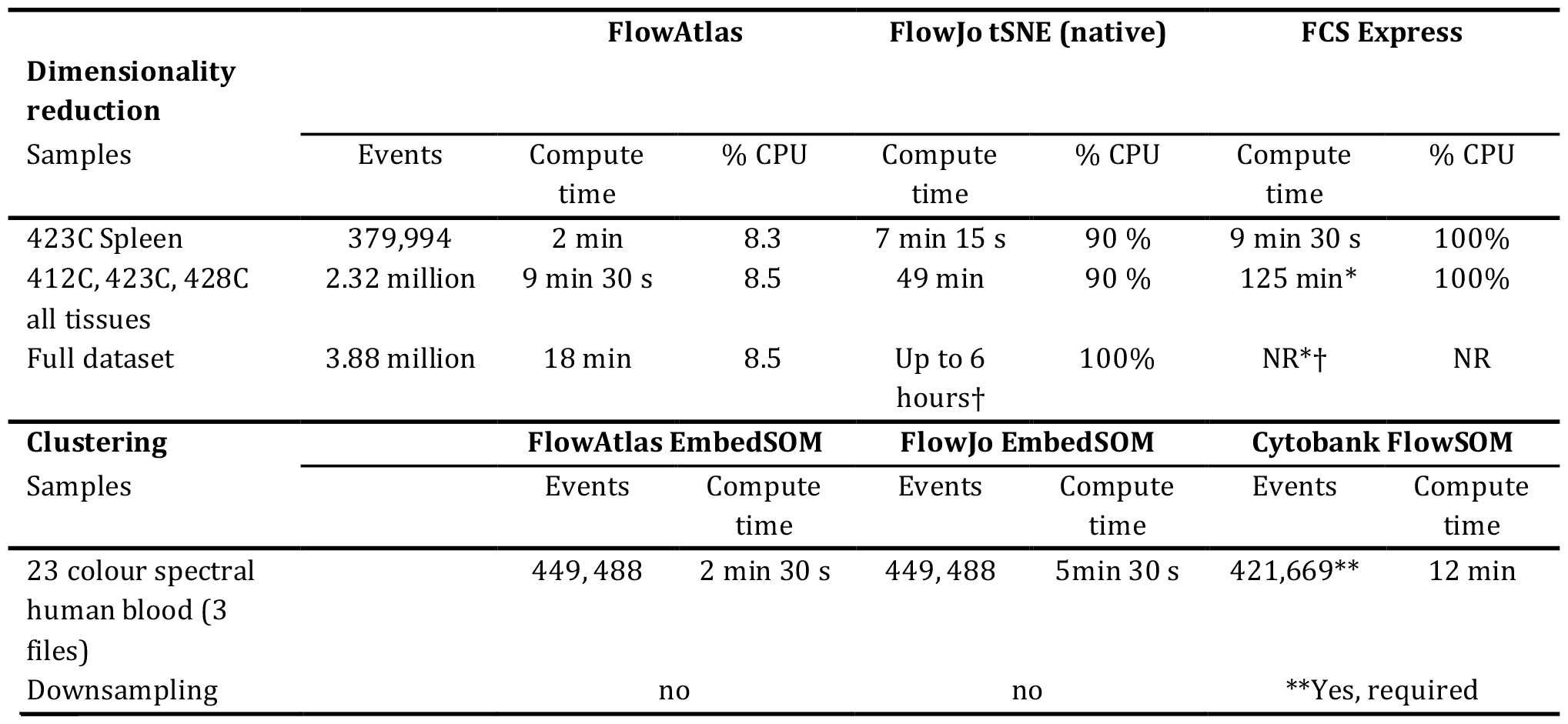
Best computational times with FlowAtlas and other high-dimensional data processing tools. CPU usage and time required by FlowAtlas, FlowJo, and FCS Express to perform dimensionality reduction on a Dell XPS15 9570 laptop with 32Gb RAM, i7-8750H CPU 2.20GHz processor. FlowJo version 10.8.1 using native tSNE tool; FCS Express version 7.18.0025. opt-tSNE settings in both platforms: all fluorescence channels, perplexity 30, iterations 1000, learning rate (eta): automatic; KNN algorithm: ANNOY, with Barnes-hut approximation (=0.5). In FlowJo and FCSExpress, different embedding topographies were produced for each sample unless samples were concatenated prior to DR. Samples stained with different panels cannot be concatenated. Times represent best results from 2-3 independent attempts. NR= not run. *Software became unresponsive on 2 of 3 previous trials. †Different panels cannot be merged, so embedding geometry varies by file. ** Downsampling required. Computation time for clustering of the indicated number of events from a publicly available spectral dataset in FlowAtlas, FlowJo, and Cytobank. The dataset is from Cytobank experiment number 191382. Time in Cytobank excludes the DR step. CPU usage is not reported for clustering analysis since it is not relevant to the cloud-based Cytobank platform.

Additionally, we compared the performance of FlowAtlas against two other non-command line clustering tools: the EmbedSOM clustering algorithm (v2.1.7) implemented as a plugin in FlowJo; and the FlowSOM algorithm implemented in the popular subscription-based cloud analysis platform Cytobank. For this test, we utilised a spectral cytometry dataset of whole human blood, which is publicly available as a demonstration experiment in Cytobank repository [10]. This dataset contains whole peripheral blood samples in 3 FCS files (23 fluorescence parameters, 512,000 events). The published data were fully unmixed and compensated; we cleaned them of debris based on scatter parameters prior to analysis, leaving 449,488 events. In FlowJo (v10.8.1), we recreated the basic gating strategy demonstrated in the Cytobank analysis to identify large major cell populations including granulocytes, B-cells, T-cells and NK cells (**Supplementary Figure 2**). We then subjected the total single cell events to DR and clustering in FlowAtlas, according to the procedure described in “Recommended FlowAtlas workflow: iterative interactive cell population discovery integrated with FlowJo“. In parallel, we replicated the demonstrated DR analysis in Cytobank (FlowSOM-on-viSNE, consensus clustering, 23 clustering parameters, without normalisation, 20 metaclusters and 100 clusters, seed 770593711). Analysis in Cytobank recommended downsampling to 420,000 events by equal random sampling (actual number of sampled events= 421,669). Lastly, we subjected the same cleaned FCS files to EmbedSOM clustering in FlowJo (v10.8.1, EmbedSOM v2.1.7; k nearest neighbours = 25, SOM grid= 20 × 20).

Clustering in Cytobank was executed in 12 minutes, excluding time required for prior viSNE dimensionality reduction. Computation in FlowAtlas took 2.5min, including embedding time. Computation in FlowJo completed in 5min 30s and, as expected, it created three embeddings with different topography (one per file, since files were not concatenated prior to analysis).

### Recommended FlowAtlas workflow: iterative interactive cell population discovery integrated with FlowJo

A typical analysis workflow using FlowAtlas concurrently with FlowJo is described in **Figure 1**. We recommend to quality control raw FCS files and remove anomalous events using dedicated tools such as FlowAI [11] or FlowCut [12]. The cleaned files should then be opened in FlowJo where the compensation accuracy is verified and live, single cells are gated and exported as new FCS files. If merging of datasets from different experiments is required for the analysis, it is recommended to consider batch-correcting the data using tools such as cyCombine [13] or CytoNorm [14] before proceeding to FlowAtlas. Following these pre-processing steps (**Figure 1** a, b-Step1), the dataset is opened in a new FlowJo workspace and antibody labels are assigned to fluorescence channels (**Figure 1** b-Step 2). Resolving naming discrepancies between channels of non-identical panels, as shown in our example, is critical because, to perform panel merging, FlowAtlas uses user-specified channel labels. FlowAtlas defaults to native fluorescence detector names when labels are not provided, which will prevent the panel merge.

Next, panel-specific gating hierarchy is created in FlowJo to define known populations of interest across all datasets (**Figure 1** b, Step-3 and **Figure 2**). This is a user-supervised population-defining step and initial annotations typically represent large populations, such as naïve/memory B-cells, or CD4/CD8 memory cells. Biexponential transformations can be applied to each channel in FlowJo, visually selecting the most appropriate co-factor for each parameter in the dataset. FlowAtlas parses the biexponential transformation directly from FlowJo for each channel, enabling the user to set optimal population separation. This in turn has been shown to dictate the quality of dimensionality reduction and metaclustering [15].

**Figure 2.**
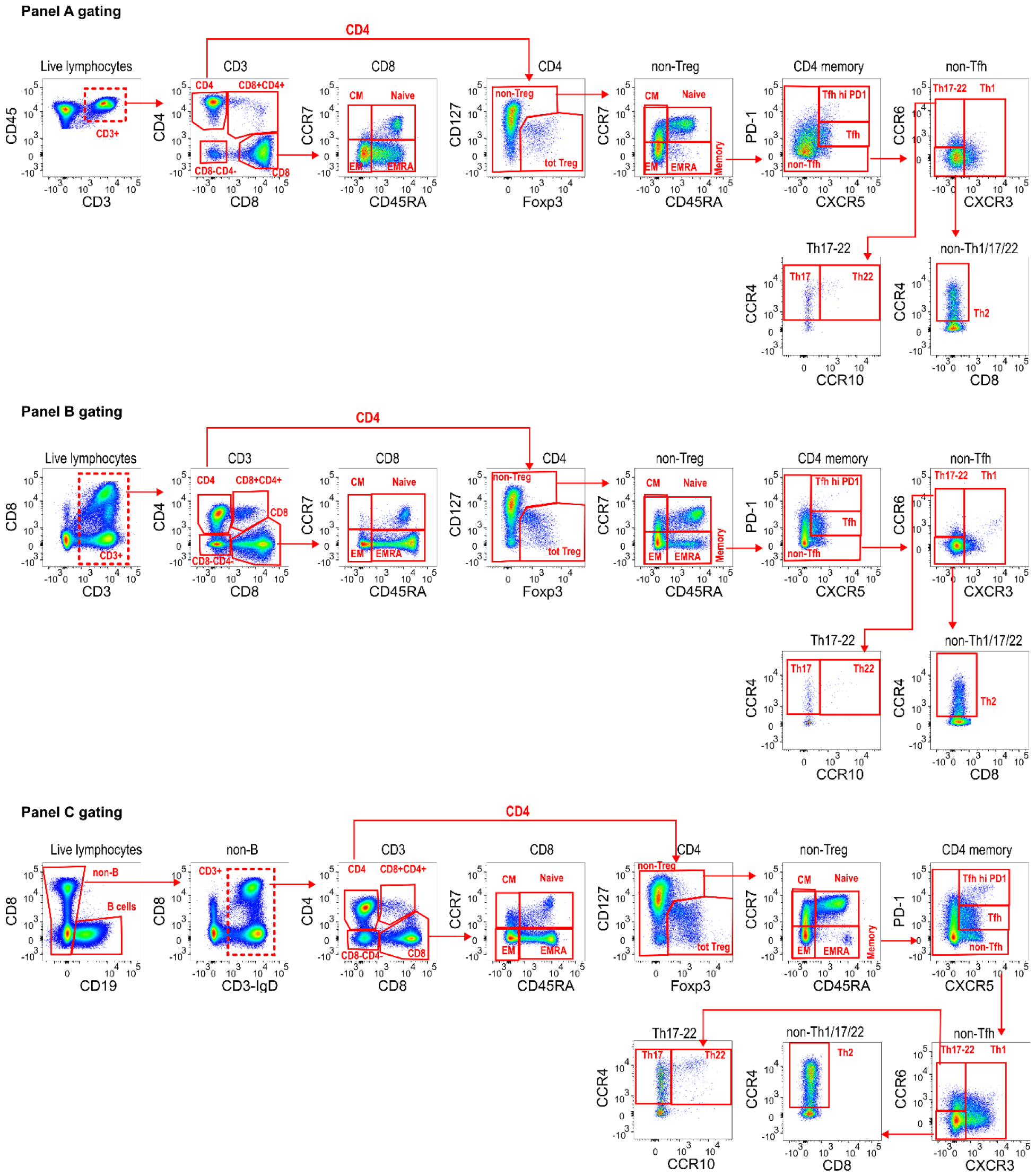
Panel-specific gating strategies created in FlowJo. For downstream DR analysis in FlowAtlas, we exported only live single T-cell events from each panel, indicated with a dashed line gate (panel A: Live CD3+ CD45+; panel B: Live CD3+; panel C: Live CD19-CD3+ events). Compensated parameters were exported, **excluding** CD45, CD19, Viability stain, FSC and SSC. Downstream gating for main population identification in FlowAtlas is shown. All channels have been biexponentially transformed. Note that other FlowJo transformations (e.g. logarithmic, ArcSinh) are not compatible with FlowAtlas.

Matching populations, irrespective of panel, are assigned the same annotation to enable cross-dataset pooling in FlowAtlas and analysis. Cells that fall outside of FlowJo-defined gates are auto-annotated as “Unlabelled” by FlowAtlas and can still be explored. Finally, to facilitate data exploration, samples are grouped by conditions enabling FlowAtlas to filter and colour-code embedded events (**Figure 1** b-Step 4). For our analysis, samples were grouped by donors and tissues. The FlowJo workspace file is then imported into FlowAtlas, which triggers panel merging, calculation of the embedding and launches an interactive browser interface (**Figure 1**c).

The user interface displays the embedding map, which can be zoomed and panned, and is rendered efficiently using OpenLayers [3].

The left-hand panel menu was designed with D3.js [4] and has four tabs: “Annotations”, “Expression”, “Frequency” and “Settings”. The “Annotations” tab enables cell filtering and re-colouring by population, condition, or by heat-map of marker expression. The filters can also be renamed or re-ordered here by dragging-and-dropping. The “Expression” tab has a polygon tool that enables drawing of multiple ROIs directly in the embedding to produce overlaid violin plots (**Figure 1** c, **Figure 3**c, **Supplementary Figure 3** and **Supplementary Figure 4**) that reveal differences in marker expression thus enabling rapid identification of clusters with unique signatures. In the “Frequency” tab frequency box plots can be generated with a few clicks (**Figure 3**a and **Supplementary Figure 5**) showing frequencies of selected populations relative to their sum or any other population. Box plot marker colours and categories displayed on the x-axis are defined by filter selections in the “Annotations’’ tab. These features enable “on-the-fly”, intuitive exploration and analysis of complex datasets. All figures can be exported as publication -quality scalable vector graphics (SVG).

**Figure 3.**
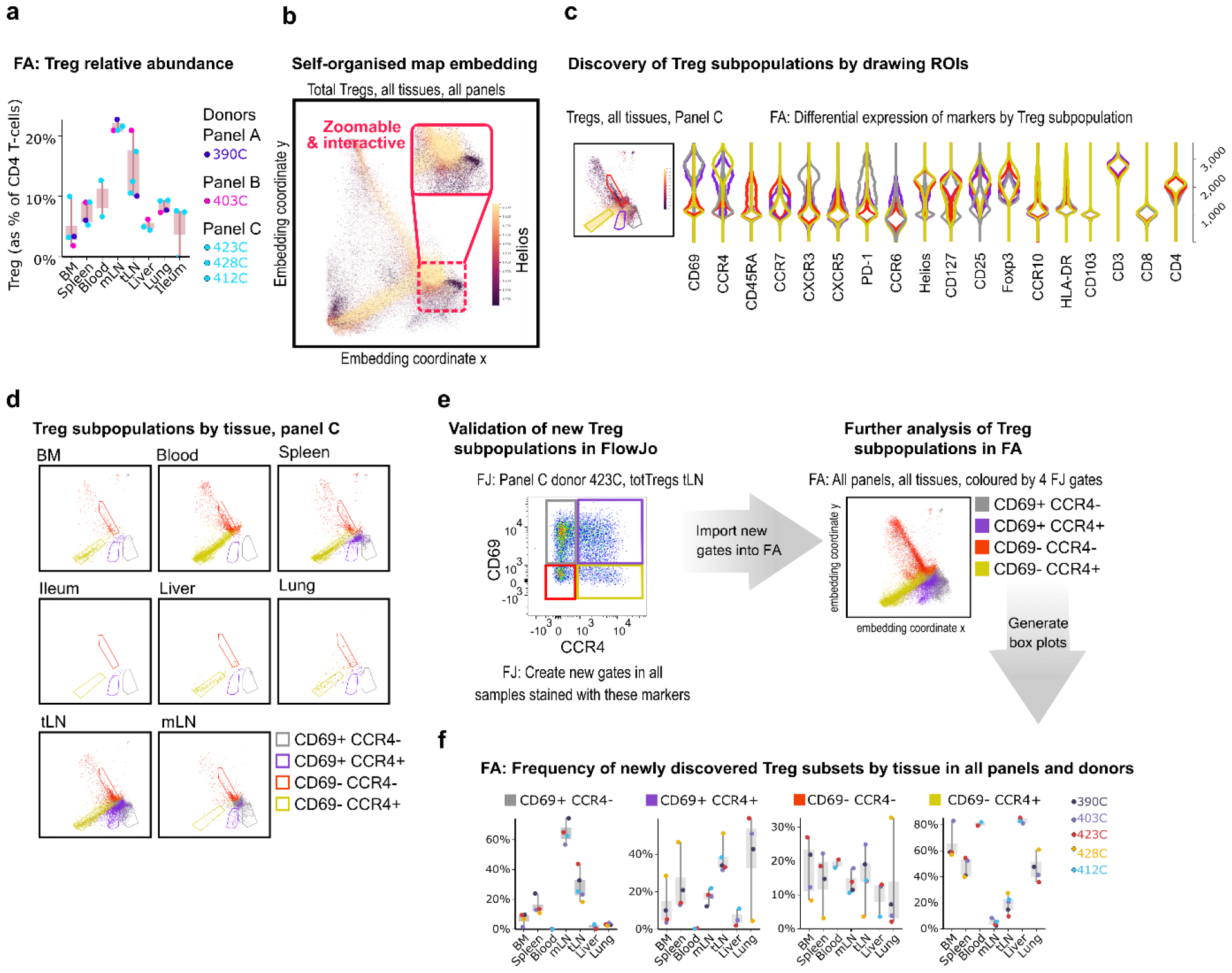
Treg subpopulation discovery in FlowAtlas. **a**, Relative abundance of Tregs by donor and tissue calculated as % of total CD4^+^ T-cells. **b**, Self-organised map embedding of Tregs from all tissues, all donors and all panels, coloured by HELIOS expression. **c**, Marker expression in four ROIs of the composite Treg embedding of all tissues stained with panel C; inset shows Tregs from all tissues stained with panel C, coloured by CCR4 expression. **d**, ROI population distributions by filtered by individual tissue. **e**, Validation and creation of new Treg sub-gates for the four ROIs in FlowJo. Gates should be created in all samples that contain the markers of interest, regardless of panel, at equivalent levels in the gating tree hierarchy (in this case, total Tregs as the parent gate). The new gates can then be opened and explored in FlowAtlas, as shown-Treg embedding re-coloured by the newly annotated Treg populations. **f**, Frequencies of the newly identified Treg subpopulations across tissues and donors. BM - bone marrow; mLN - mesenteric lymph nodes; tLN - thoracic lymph nodes; ROI - region of interest; FJ - FlowJo; FA - FlowAtlas.

Once unique subpopulations have been identified, they can be validated in FlowJo with targeted two-parameter plots and new gates created to be read by FlowAtlas at rerun. This “iterative discovery loop” substantially simplifies discovery.

Embedding is performed only once when the workspace file is first imported and is stored in a cache file with a “.som” extension, allowing users to return to their analysis quickly. The embedding can also be re-calculated to change cluster geometry. Sharing the “.som” cache file together with the FlowJo workspace and FCS files enables collaboration, allowing colleagues to work on the same embedding map.

Hereafter, we demonstrate the capabilities of FlowAtlas using our novel conventional cytometry dataset of multi-donor multi-tissue derived immune cells. Utilisation of FlowAtlas for analysis of spectral and CyTOF data is shown in **Figure 4** and **Supplementary Figure 6** respectively.

**Figure 4.**
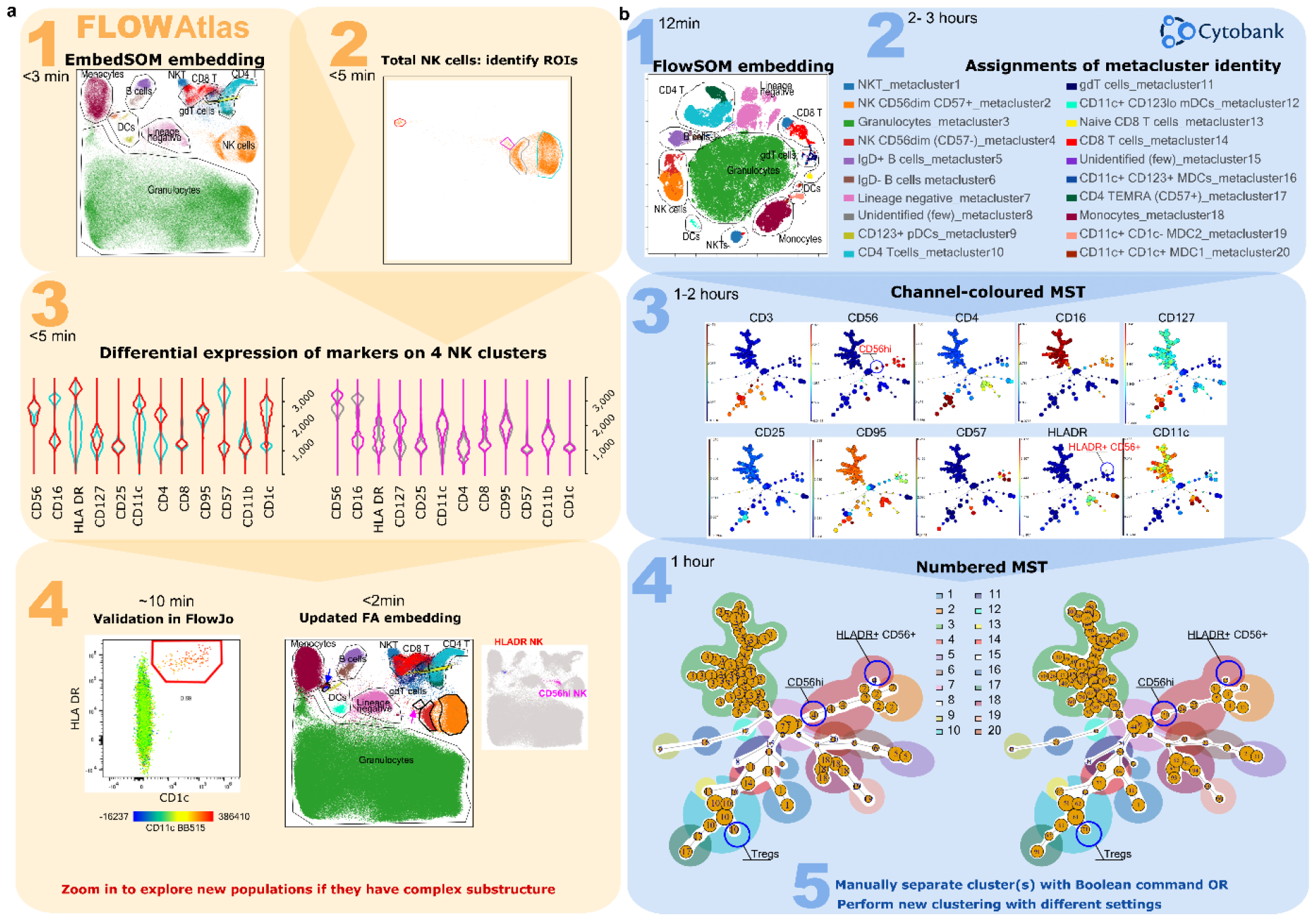
Comparison of workflow for the detection of rare cell subsets in FlowAtlas. (a) and Cytobank (b) using a published spectral cytometry 23-colour dataset of whole human blood. In FlowAtlas, embedding the data is quick, and basic populations are created using the familiar workflow of FlowJo (Step 1a). The eye is immediately drawn to heterogeneity in the embedding, for example in the NK cell population (Step 2a). A small cluster of NK cells (magenta ROI and violins) is close to the myeloid populations, and expresses HLA-DR, CD11c and CD1c (Step 3a). Checking that it exists in FlowJo (Step 4a) is easy- and it is quickly added to the updated FlowAtlas embedding. The user can now also zoom in on three other larger NK subsets (CD56bright, CD57+ CD56dim and CD57-CD56dim). The equivalent workflow in Cytobank is as follows: after a relatively fast embedding of the data (step 1b), the user needs to identify each of the 20 metacluster identities first (step 2b), using heatmaps and violin plots of marker expression (not shown). Rare populations may not have segregated. They can be discovered by examining the MST, coloured by channel (Step 3b and 4b)-looking for heterogeneity in a metacluster. For example, metacluster 4 contains cluster 15, which appears to express HLA-DR and CD56.The user can either re-run the analysis, or create new metaclusters by manually combining cluster numbers (Step 5b). Equivalent major cell populations are coloured identically in the two embeddings and in the minimum-spanning trees (in Step 4b); ROI colour in FlowAtlas matches the corresponding violin plots.

## Demonstrating the utility of FlowAtlas

### Example cell population exploration

Our dataset consists of 32 files of tissue-derived immune cells obtained from 5 deceased transplant organ donors (**Supplementary Table 1**), stained with 3 different panels (**Supplementary Table 2**). The data were pre-processed in FlowJo to remove anomalous events, debris and aggregates; compensation was checked; and live, single T cells were exported as new FCS files for downstream analysis. These files were imported into a new FlowJo workspace and each channel was biexponentially transformed, basic populations were gated (**Figure 2**), and samples were grouped by donor ID and source tissue. Next, DR and clustering were performed in FlowAtlas. After generating relative abundance boxplots of the major lymphocyte populations in our dataset (**Supplementary Figure 5**), we elected to zoom into the CD4 regulatory T-cell (Treg) compartment, defined as CD3^+^CD4^+^CD127^-/lo^FOXP3^+^ cells, as an exemplar.

As a proportion of all CD4^+^ T-cells,Tregs were demonstrated to be enriched in lymph nodes, particularly mesenteric lymph nodes where they accounted for more than 20% of CD4 T-cells in all studied donors (**Figure 3** a).

The embedding of Tregs for Panel C donors, recoloured by the expression of the transcription factor HELIOS (**Figure 3** b), revealed the presence of HELIOS^+^ and HELIOS^-^ subpopulations as expected [16,17], with additional subcluster structures. Next, we filtered the embedding by panel C samples and used it to explore Treg subcluster characteristics further. We coloured embedded events by tissue of origin and drew ROIs around four main subclusters seen in the embedding (**Figure 3** c). Auto-generated violin plots quickly allowed us to observe differences in expression of CD45RA, CCR7, CCR4 and CD69 between these subclusters, with the red ROI having a naive phenotype (CD45RA^+^CCR7^+^) and lacking CCR4 and CD69 expression, while yellow, grey and violet ROIs showed characteristics of memory subsets (CD45RA^-/lo^CCR7^-^) with and without CD69 and CCR4 expression. Filtering the embedding by tissue with the above ROIs superimposed (**Figure 3** d), revealed tissue-specific enrichment patterns; for example, CD69^+^ subsets were largely absent from blood, consistent with the role of CD69 in promoting tissue retention [18] [19] [20], whereas liver, lung, and thoracic lymph nodes contained a high proportion of Tregs expressing the chemokine receptor CCR4^+^ (with or without CD69 co-expression).

CCR4 has been implicated in T-cell trafficking to the lung [21], and in the infiltration of Tregs into tumours [22]. Next, we validated the presence of these four Treg subsets in FlowJo (**Figure 3** e) and created new gates using CCR4 and CD69-now in all samples stained with these markers, irrespective of panel-for further exploration in FlowAtlas. Returning to FlowAtlas, we re-coloured the Treg embedding by these newly annotated subsets and generated frequency box plots (**Figure 3** f), which further highlighted tissue-specific expression patterns.

FlowAtlas allowed us to obtain deep insights into the Treg population rapidly and intuitively. Therefore, we applied a similar analysis strategy to CD4^+^ Th1 and CD8^+^ memory cells, producing further data in a matter of minutes (**Supplementary Figure 3** and **Supplementary Figure 4**). This contrasts with analysis solely performed within FlowJo, where the computation of our full dataset embedding of 3.88 million events using tSNE would have been prohibitively slow (6h, see **Table 1** for comparison of performance) and assessing all possible combinations of markers using two-dimensional plots would have been a laborious process.

Although EmbedSOM is now implemented as a FlowJo plugin, downstream exploration of the resulting embedding still relies on classic 2-way scatter plots and cannot be zoomed or easily filtered by custom conditions. Furthermore, preserving a consistent topography across samples requires either file concatenation, or clustering by FlowSOM first-both of which impose that all samples are stained with the same panel. FlowAtlas overcomes these barriers, making the embedded data interactive, and patterns within it-quickly visible.

### Detection of rare cell subsets using FlowAtlas

Current DR computational pipelines reduce computation time by downsampling large datasets using random uniform sampling, which may not optimally reflect the distribution of the original data [23]. Rare cell subsets may be missed by data downsampling and underfitting in existing unsupervised clustering approaches. Since FlowAtlas does not downsample, it potentially circumvents this problem.

Accordingly, we tested the ability of FlowAtlas to discover novel rare cell populations in the above-mentioned 23-parameter spectral cytometry dataset of whole human blood [10]. As described, we performed the analysis in FlowAtlas and then replicated the example analysis demonstrated in Cytobank from curated experiment number 191382. The gating strategy for this dataset is shown in **Supplementary Figure 2**. Using FlowAtlas, we identified a subset of HLA-DR^+^ NK cells, comprising only 0.69% of total NK cells in under 30 min (**Figure 4**, steps 1a-4a). The same population was not resolved as a separate metacluster in Cytobank FlowSOM-on-viSNE analysis at the implemented settings (**Figure 4**, steps 1b and 2b). Furthermore, CD56^bright^ NK cells, which are well known to be phenotypically and functionally distinct [24], also did not segregate from the main NK cell population at these analysis settings. In order to find the missing HLA-DR^+^ CD56^+^ subpopulation in Cytobank, it was necessary to review the 10 individual clusters comprising CD56^+^ events in the minimum spanning tree (MST), coloured by each channel median fluorescence intensity (MFI), which was a time-consuming process. We noted that cluster 15 within metacluster 4 was located away from the main metacluster 4 nodes and that it contained a small subset of HLA-DR^+^ CD56^+^ NK cells (**Figure 4**, step 3b and 4b). These may be the equivalent population to the cells discovered in FlowAtlas. We verified that the other 9 neighbouring NK-cell clusters did not contain this population, by examining scatter plots of their key identifying markers (HLA-DR, CD11c) versus cluster number (not shown). Finally, we isolated the subpopulation manually based on its cluster number. This process took several hours and was informed by our prior identification of this population in FlowAtlas.

Resolution of other rare populations would potentially require each of the 100 clusters in the MST to be individually examined, as above. Once discovered, a rare subpopulation would either need to be manually isolated by combining the clusters that contain it with Boolean commands, or a new clustering would need to be undertaken with optimised settings or starting with a purer cell population. By contrast, FlowAtlas allows the user to simply zoom in on the existing embedding to study the substructure of clusters without needing to re-embed the data.

### FlowAtlas can integrate multiple flow cytometry panels, but protocol-driven experiment harmonisation remains critical

Due to an evolution of panel design, our tissue-derived immune cell dataset consisted of 3 different panels. Most existing computational tools require the concatenation of files prior to analysis, which is impossible when different markers have been assigned to the same channel. This would typically cause researchers to exclude precious data that they cannot integrate.

FlowAtlas enables data re-use and concomitant analysis of datasets acquired with non-identical antibody panels by imputing missing values using random sampling with replacement before DR. Algorithmic bias is prevented by excluding imputed values from the embedding visualisation or any downstream analyses.

To demonstrate the capability to merge panels, we acquired 2 healthy control blood samples and stained them with the 3 panels previously used in our main tissue-derived dataset. We integrated the 6 new FCS files (1.28 million live single T-cell events) into the existing embedding of tissue-derived immune cells. The use of the same two donors eliminated any biological variation, enabling us to isolate the effect of panel differences within the healthy control group.

We filtered the embedded data by “healthy control”, then coloured the embedding by panel and inspected differences in cluster position, geometry and marker mean fluorescence intensity (MFI). Violin plots revealed variation in marker expression due to panel design differences, e.g. lower CD4 MFI for panels B and C compared to panel A, due to use of different fluorochromes (see **Supplementary Table 2**). The overall embedding geometry was highly conserved across the three panels (**Figure 5**a).

**Figure 5.**
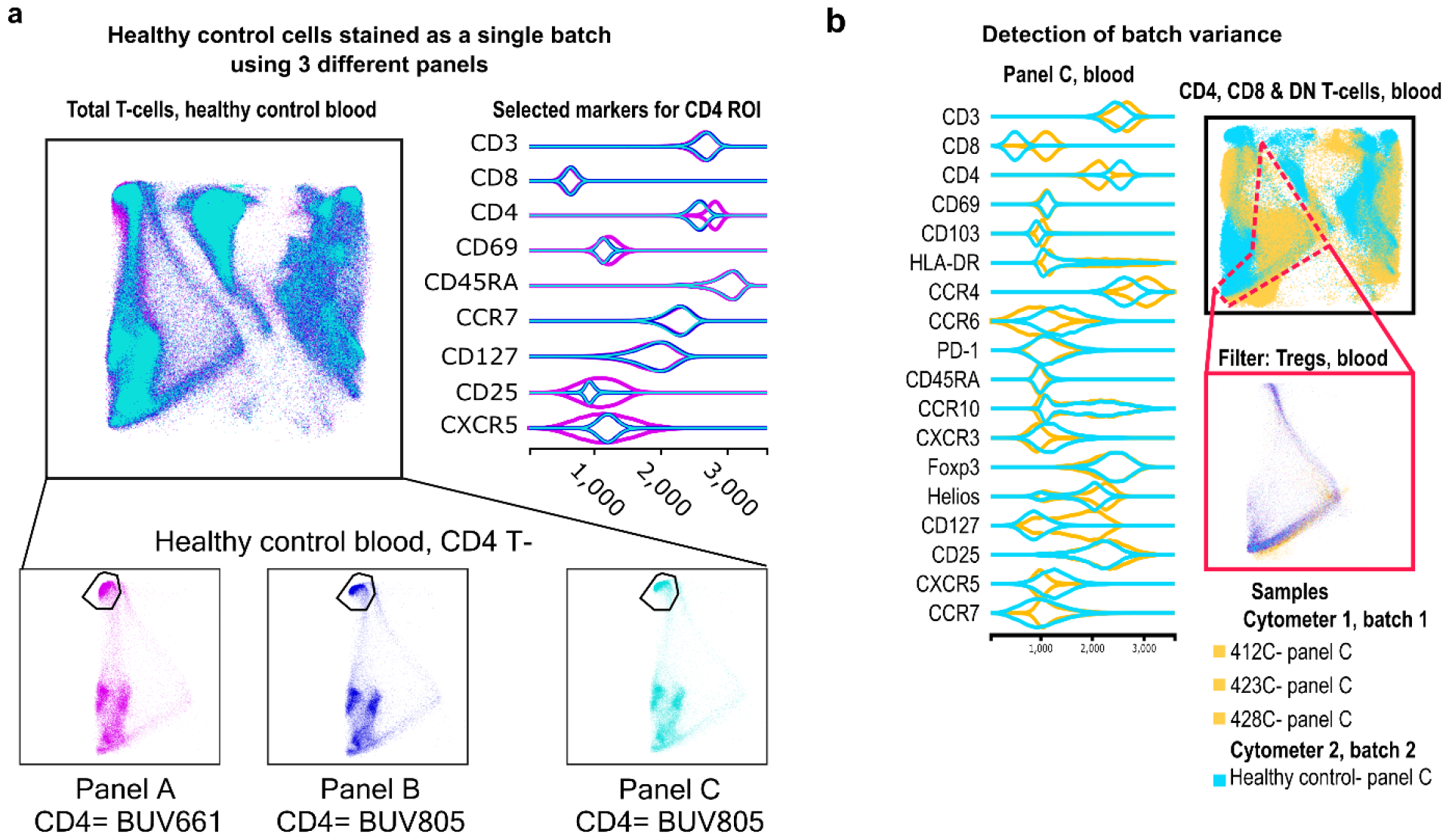
Merging of panels and detection of batch variance. a, Two healthy control donors were stained with our 3 panels as one batch, and data were processed in FlowAtlas as recommended. Events are coloured by panel and show minimum differences in population geometry, driven by our choice of CD4 fluorochrome (BUV661 on panel A, BUV805 on panels B and C). Spread from BUV661 into CD25 -APC and CXCR5-APCR700 is visible in panel A in violin plots. The three panels integrate well without normalisation.b) Blood samples stained with panel C are shown as embedding and violin plots (yellow= deceased organ donor blood, processed ex vivo is designated “batch 1”; cyan= healthy control blood, processed after cryopreservation and designated “batch 2”). FlowAtlas has successfully merged the panels, but the topography is very different between the two batches. This likely represents a mixture of biological differences, and in large part-batch differences due to different sample handling and cytometer configurations.

Next, we filtered samples stained with panel C, displayed only blood-derived cells, and coloured the samples by batch (healthy controls vs deceased organ donors, **Figure 5**b). We noted significant qualitative differences in the embedding geometry for these two sets of samples. The resulting violin plots showed differences in several chemokine receptors, CD127, CD4 and CD8. Although biological differences between healthy and deceased donor blood could account for this observation, the magnitude of the differences strongly suggested an additional batch effect, potentially due to the fact that, unlike the tissue-derived dataset, healthy PBMCs had been cryopreserved and acquired on a cytometer with a different optical configuration (See Methods and **Supplementary Table 4**). This was unsurprising, but important to highlight, given that FlowAtlas does not perform MFI normalisation.

We have noted that panels with very few shared markers and/or fluorochromes can still be processed but in this case, equivalent populations will likely fail to co-localise adequately due to a lack of common landmarks (**Supplementary Figure 7**).

In summary, FlowAtlas is relatively robust at handling samples with moderately different panels where marker MFIs have not been normalised, but optimum co-localisation of equivalent populations requires batch-normalisation at pre-processing, and that panel differences are minimal.

## Discussion

FlowAtlas is a novel open-source data exploration tool, which combines the computational power of the GigaSOM library and Julia programming language with the widely used software FlowJo, expanding its capabilities in a completely graphical, fast, user-friendly interface. This approach removes all entry barriers imposed by command-line analysis pipelines that currently hold many users back from taking advantage of powerful computational tools. FlowAtlas brings a new iterative analysis concept to biomedical scientists by linking the familiar FlowJo workflow with a high-performance machine learning framework. FlowAtlas allows rapid computation of millions of high-dimensional events without the need for down-sampling. The highly interactive embedding enables zooming and intuitive exploration of population substructure, considerably speeding up population discovery. Missing-data handling methods enable concomitant analysis of datasets with non-identical panel designs or markers.

Importantly, FlowAtlas does not incorporate batch normalisation, and, to prevent algorithmic bias, does not display imputed values in the embedding. Here, we briefly discuss the rationale behind our design decision.

Firstly, we designed FlowAtlas without a data normalisation step so that users can select the most appropriate method for eliminating technical variability for their specific experimental context at the data pre-processing step.

Best practice currently relies on inter-laboratory protocol harmonisation through the use of standardised antibody cocktails, identical staining procedures, calibration of cytometers using fluorescence standards or Application Settings [25] and internal biological “anchor” controls stained with each batch of samples. Anchor controls enable batch correction using pipelines such as swiftReg in R [26] and CytoNorm [14]. The latter is available as a FlowJo plugin and circumvents the need for coding expertise. These protocol-based approaches, which we used to acquire our tissue-derived immune cell dataset, would likely best suit the primary target user demographic of FlowAtlas.

In the absence of internal anchor controls, the currently available computational methods of batch correction mostly require considerable command-line competence. For example, GaussNorm (in R) aligns cellular landmarks (positive and negative population peaks) across samples [27]. This approach may eliminate biologically relevant MFI differences and is only suitable when population frequency is the variable of interest. Powerful batch correction tools rooted in single-cell genomics packages are now finding application in flow and mass cytometry, e.g. Seurat (in R) [28] and Pytometry (in Python) [29]. The stringency of batch effect removal versus biological effect preservation varies widely between these methods [30], so the optimum analysis pipeline may vary between datasets. These pipelines were originally developed to handle very high-dimensional data with thousands of observations per parameter and high batch variability (e.g. different technology platforms). Datasets with very few observations per sample, in which batch effects are driven by few parameters, as was the case in our tissue-derived dataset, may not be amenable to these correction methods.

Secondly, panel merging and missing-data handling methods in FlowAtlas ensure it is relatively robust to moderate panel differences, enabling dataset integration in selected circumstances. We substituted some markers in our panels with completely spectrally unique fluorochromes and demonstrated that FlowAtlas can preserve the embedding geometry under the tested conditions. Nevertheless, panels with little overlap in markers or fluorochromes are unlikely to integrate successfully. Where multiple markers differ, users are advised to test the effectiveness of panel integration by staining a single donor sample with their panels of interest and assessing the resulting embedding geometry. Tools have been developed, which aim to combine panels through marker imputation, e.g. CytoBackBone [31], CyTOFMerge [32], Infinicyt (Cytognos, BD) and CyCombine [13]. Nevertheless, we chose not to display imputed values in the FlowAtlas embedding to protect against bias. A critical assessment of these methods has recently reported relatively poor approximation of known expression values [33], justifying our decision.

In conclusion, FlowAtlas is a novel data exploration tool, which leverages advanced machine learning methods, rapid computational speed, and a near-complete lack of a user learning curve before data exploration can commence. The highly interactive and intuitive workflow eliminates the need for command-line coding and brings high-dimensional data exploration and population discovery to the non-bioinformatician biologist.

## Methods

### Ethical statement

All work was completed under ethically approved studies. Healthy human PBMCs were isolated from volunteers having given informed consent under CAMSAFE (REC-11/33/0007). All deceased organ donor tissue samples were collected via the Cambridge Biorepository for Translational Medicine under Research Ethics Committee approval 15/EE/0152. In addition, two donor-matched blood samples were collected prior to withdrawal of life support, under Ethics Committee approval 97/290.

### Tissue acquisition and dissociation, and preparation of healthy control PBMCs

Tissue was obtained from five deceased organ donors following circulatory death. Donor metadata is given in **Supplementary Table 1**. Briefly, following cessation of circulation, human donor organs were perfused in situ with cold organ preservation solution and cooled with topical application of ice. Samples for the study were obtained within 60 minutes of cessation of circulation and placed in University of Wisconsin organ preservation solution for transport at 4°C to the laboratory. Lung and liver samples were obtained from the left lower lobe of the lung and the right lobe of the liver. In addition, two donor-matched blood samples were collected prior to withdrawal of life support (under REC approval 97/290). To minimise the possibility of processing-dependent differences in cell surface marker expression, all samples, including blood, were processed using enzymatic digestion protocol. Briefly, solid tissues were weighed, transferred into 10cm tissue culture dishes, and cut into small pieces. Up to 5g of tissue was then transferred into a GentleMACS C tube (Miltenyi Biotec) prefilled with 5mL of dissociation media composed of X-VIVO15 with 0.13U/mL Liberase TL (Roche), 10U/mL Benzonase nuclease (Millipore/Merck), 2% (v/v) heat-inactivated foetal bovine serum (FBS, Gibco), penicillin (100 U/ml, Sigma-Aldrich), streptomycin (0.1 mg/ml, Sigma-Aldrich), and 10mM HEPES (Sigma Aldrich). The samples were then homogenised using a GentleMACS Octo dissociator (Miltenyi Biotec) running a protocol that provided gradual ramping up of homogenisation speed and two 15 - minute heating/mixing steps at 37°C. Digested tissue was passed through a 70μm MACS Smartstrainer (Miltenyi Biotec) and the flow-through was first washed with X-VIVO15 supplemented with 2 mM EDTA and then with PBS. Mononuclear cells were enriched by Ficoll-Paque (GE Healthcare) density centrifugation according to the manufacturer’s instructions. Following density centrifugation, mononuclear layer was collected, washed once with PBS and the cell pellet was resuspended in FACS buffer (PBS, 2.5% FBS). Bone marrow aspirates and peripheral blood samples were first subjected to Ficoll-Paque density centrifugation, according to manufacturer’s instructions, the mononuclear layer was then collected, washed with PBS and cells were treated with the same dissociation media as solid tissues for 30 min at 37°C prior to washing and resuspension in FACS buffer.

Healthy control PBMCs were prepared by Ficoll-gradient centrifugation and cryopreserved in cell freezing medium (Sigma) containing 10% DMSO for future use.

### Flow cytometry of tissue-derived mononuclear cells

Depending on the cell yield, up to 1x10^6^ mononuclear cells/tissue were stained with antibodies shown in **Supplementary Table 3**. Not all donors were stained with the same panel. To expand the total number of markers, sentinel panel design was implemented where CD3 and IgD were detected with antibodies conjugated to BUV395 and FOXP3 and IgM were detected with antibodies conjugated to PE in some donors. Refer to **Supplementary Table 2** for details. Single cell suspensions were washed once in PBS, transferred into 96 v-bottom plate and stained with Zombie UV viability dye for 30 min at 4°C followed by a wash with FACS buffer. Cell pellets were resuspended in 50μl FACS buffer with Human FcR block (BD Biosciences) and incubated for 10 min at 4°C. Next, cells were pelleted, excess buffer removed and 100μl of antibody master mix composed of cell-surface antibody cocktail (see **Supplementary Table 2**), BV buffer (BD) and True-Stain Monocyte Blocker (Biolegend) and incubated for 1h at 4°C. Following incubation, cells were washed three times in PBS and prepared for intracellular staining using transcription factor fixation/permeabilisation kit (eBioscience) according to the manufacturer’s instructions. Following intracellular staining, cells were resuspended in PBS and analysed on BD FACSymphony A3 cell analyser within 10 hours.

### Flow cytometry of healthy PBMCs

In contrast to tissue-derived samples, which were processed *ex vivo*, healthy PBMC samples were thawed in X-VIVO15/10% FCS at room temperature and stained according to the procedure above. Analysis was performed on a BD FACSymphony A5 cell analyser within 10 hours. The optical configuration of the two cytometers used in this study is shown in **Supplementary Table 4**. The cytometers were not cross-calibrated for comparable measurement of MFI, but each underwent individual CS&T bead quality control before sample acquisition.

### Computational methods and step-by-step instructions for FlowAtlas use

#### Flow cytometry data pre-processing

Raw FCS data were cleaned using FlowAI [11] to remove acquisition anomalies. High-quality files were saved and imported into FlowJo for data pre-processing. In this step, compensation matrices were curated; aggregates and dead cells were gated out; and remaining cells were gated on lymphocytes and T-cells (see **Figure 2**). The live T-cell gate of each sample was exported as a new FCS file containing only compensated fluorescence channels. The pre-processed files were then opened in a new FlowJo workspace, where antibody labels were assigned to all fluorescence channels.

Compensated parameters were exported for live single aggregate-free T-cells from all panels for dimensionality reduction, since we found that this produced optimal geometry for analysing the T-cell population, which was of particular interest. Therefore, we labelled the PE channel as FOXP3 in all files. Also, CD4 was used on either BUV661 (Panel A) or BUV805 (Panels B and C).

It is possible to subject the entire Live Singlet aggregate-free population to DR analysis if desired. In this case, CD19^+^ events would only be identifiable in samples labelled with panel C; equivalent cells would appear as “Ungated” in other panels. For this workflow, we would recommend labelling the PE channel in all datasets as FOXP3-IgM; this would display both PE-labelled markers on a single violin plot; separate downstream differential expression analysis of each marker is made possible by filtering events by cell type (B-cell or T-cell) in FlowAtlas.

#### FlowAtlas code availability

The code for FlowAtlas is open-source and is available at our GitHub repository: https://github.com/gszep/FlowAtlas.jl.git

#### Installation and Loading of FlowAtlas

FlowAtlas is compatible with FlowJo version 10.8.1.

FlowAtlas requires Julia language, which is easily installed on any operating system by downloading an installer available here: https://julialang.org/downloads and following the on-screen instructions. Tick the option to add Julia to PATH environment when prompted.

Once Julia is installed, FlowAtlas can be installed and run in three lines of code as follows:

1. Windows: open Run (Windows Key + R), type **cmd** and hit enter. MacOS: open command prompt (Cmd Key + Space), type **terminal** and hit enter. This will launch Windows/MacOS command prompt.
2. In the prompt type **Julia** and hit enter. This will launch the Julia environment.
3. Type ] and the prompt will change to display that package manager is now active.
4. Type **add FlowAtlas** and hit enter. This will download and install FlowAtlas.jl. Once installation is complete, you can close the command prompt window.

To start using FlowAtlas, navigate to the folder containing your pre-processed FCS files (make sure that the FlowJo workspace file is there as well) and launch command prompt: in Windows by typing **cmd** in the File Explorer address bar (where file path is usually displayed) and hitting enter or in MacOS launch terminal and navigate to the folder by typing cd followed by the folder path. In the prompt, type **Julia** and hit enter to start it, then type **using FlowAtlas** and hit enter. Once FlowAtlas is loaded, type **FlowAtlas**.**run(“workspace**.**wsp”; files=“*/\***.**fcs”)** where **workspace**.**wsp** is the name of your FlowJo analysis file with .wsp extension. Adding new files into the workspace after initial analysis will force a recalculation of the embedding.

A short video demonstrating the use of FlowAtlas can be watched here: https://www.youtube.com/watch?v=FeYrFKgP91s

## Declarations

### Data availability

All data are made available in FlowRepository http://flowrepository.org/id/FR-FCM-Z74D.

## Acknowledgements

We thank the deceased organ donors, donor families, the extended Cambridge Biorepository for Translational Medicine team, and the transplant coo rdinators for access to the tissue samples. We also thank Professor Linda Wicker (University of Oxford) for her academic insights. This work was funded in part by the Wellcome Trust (Grant number 105924/Z/14/Z; RG79413 to JLJ). For the purpose of open access, the authors have applied a CC BY public copyright licence to any Author Accepted Manuscript version arising from this submission. This work was also supported by the NIHR Cambridge Biomedical Research Centre (BRC121520014), and by the Cambridge NIHR BRC Cell Phenotyping Hub. The views expressed are those of the author(s) and not necessarily those of the NIHR or the Department of Health and Social Care. GS was supported by Microsoft Research and by the EPSRC Centre for Doctoral Training in Cross-Disciplinary Approaches to Non-Equilibrium Systems (CANES, EP/L015854/1). ZG was supported by the Wellcome Trust (Grant number 220554/Z/20/Z). HSM was supported by The Rosetrees Trust (RG82826, JS16/M589).

## Ethics

JLJ reports receiving consultancy fees and grant support from Sanofi Genzyme. All other authors declare no competing interests.

