## Supplementary Table 1 for "FlowAtlas.jl: an interactive tool bridging FlowJo with computational tools in Julia"

| Donor ID | Sex | Age | Primary cause of death | Multi trauma | Days in Hospital | BMI | CMV EBV TOXO | Smoking | Alcohol (u/day) | Antibiotics within 2 weeks of death | Steroids |
|---|---|---|---|---|---|---|---|---|---|---|---|
| 390C | F | 65-70 | ICH | ✓ | 2 | 30- 35 | +/+/- | ? | <1 | × | × |
| 403C | M | 50-55 | ICH | ✓ | 8 | 30- 35 | +/+/- | ✓ | <1 | Co,T | × |
| 423C | M | 60-65 | ICH | × | 2 | 20- 25 | -/+/- | ✓ | >9 | G,F | D |
| 412C | M | 70-75 | ICH | × | 5 | 26- 30 | -/+/+ | ✓ | <2 | A*, F, G, C, Co | P† |
| 428C | F | 55-60 | ICH | × | 3 | 20- 25 | -/+/- | ✓ | >9 | Co | × |

F= Female; M= Male; ICH= intracranial haemorrhage; CMV= cytomegalovirus; EBV= Ebstein-Barr virus; TOXO= Toxoplasmosis; Co= co-amoxiclav; A= amoxicillin; T= Tazocin; F= Flucloxacillin; G= Gentamicin; D= Dexamethasone; C= Clarithromycin; ✓= yes;×= no; ?= not known; P= Prednisolone; * pre-admission; † pre-treatment

***Supplementary Table 1 Donor characteristics***
