## Supplementary Table 2 for "FlowAtlas.jl: an interactive tool bridging FlowJo with computational tools in Julia"

| Specificity | Fluorochromes | | | |
|---|---|---|---|---|
| | Panels | A | B | C |
| | Donor(s) | 390C | 403C | 412C, 423C, 428C |
| CD45 | | BUV805 | - | - |
| CD19 | | - | - | BUV570 |
| IgM | | - | - | PE |
| IgD | | - | - | BUV395 |
| CD4 | | **BUV661** | **BUV805** | **BUV805** |
| CD3 | | BUV395 | BUV395 | BUV395 |
| CD8 | | BUV563 | BUV563 | BUV563 |
| CD69 | | BUV737 | BUV737 | BUV737 |
| CD103 | | BV421 | BV421 | BV421 |
| HLA-DR | | BV510 | BV510 | BV510 |
| CD127 | | PE-Cy7 | PE-Cy7 | PE-Cy7 |
| CCR4 | | BV605 | BV605 | BV605 |
| CCR6 | | BV650 | BV650 | BV650 |
| PD-1 | | BV711 | BV711 | BV711 |
| CD45RA | | BV786 | BV786 | BV786 |
| CCR10 | | BB515 | BB515 | BB515 |
| CXCR3 | | BB700 | BB700 | BB700 |
| CXCR5 | | APCR-700 | APCR-700 | APCR-700 |
| CCR7 | | APC-Fire750 | APC-Fire750 | APC-Fire750 |
| CD25 | | APC | APC | APC |
| FOXP3 | | PE | PE | PE |
| HELIOS | | PE-Dazzle | PE-Dazzle | PE-Dazzle |
| Live/Dead | | Zombie UV | Zombie UV | Zombie UV |

***Supplementary Table 1 Immunophenotyping panels used in this study**. Different fluorochromes were used for anti-CD4 in each panel (in bold). Not all antibodies were present in all panels. FOXP3 and IgM were both on PE in panel C.*
