## Supplementary Table 3 for "FlowAtlas.jl: an interactive tool bridging FlowJo with computational tools in Julia"

| Specificity | Fluorochrome | Clone | Source |
|---|---|---|---|
| CD3 | BUV395 | SK7 | BD |
| IgD | BUV395 | IA6-2 | BD |
| CD8 | BUV563 | RPA-T8 | BD |
| CD69 | BUV737 | FN50 | BD |
| CD4 | BUV661 | SK3 | BD |
| CD4 | BUV805 | SK3 | BD |
| CD45 | BUV805 | HI30 | BD |
| CD103 | BV421 | Ber-ACT8 | BD |
| HLA-DR | BV510 | G46-6 | BD |
| CD19 | BV570 | HIB19 | Biolegend |
| CCR4 | BV605 | L291H4 | Biolegend |
| CCR6 | BV650 | 11A9 | BD |
| PD-1 | BV711 | EH12.1 | BD |
| CD45RA | BV786 | HI100 | BD |
| CCR10 | BB515 | 1B5 | BD |
| CXCR3 | BB700 | 1C6/CXCR3 | BD |
| FOXP3 | PE | 259D/C7 | BD |
| FOXP3 | PE | PCH101 | eBioscience |
| IgM | PE | G20-127 | BD |
| HELIOS | PE-Dazzle594 | 22F6 | Biolegend |
| CD127 | PECy7 | HIL-7R-M21 | BD |
| CD25 | APC | 2A3 | BD |
| CD25 | APC | MA251 | BD |
| CXCR5 | APC-R700 | RF8B2 | BD |
| CCR7 | APC-Fire750 | G043H7 | Biolegend |
| Viability | Live/Dead UV | - | Biolegend |

***Supplementary Table 1 Details of antibodies used in this study***
