## Supplementary Table 4 for "FlowAtlas.jl: an interactive tool bridging FlowJo with computational tools in Julia"

| Laser | FACS Symphony A3 | FACS Symphony A5 | Utilised fluorochrome |
|---|---|---|---|
| Acquired samples | Tissue-derived immune cells | Healthy control PBMCs | |
| | Bandpass filter | Bandpass filter | |
| 355nm | 379 / 28 | 379 / 28 | BUV395 |
| | | 450 / 50 | - |
| | 515 / 30 | 515 / 30 | Zombie UV |
| | 560 / 40 | 580 / 20 | BUV563 |
| | | 610 / 20 | - |
| | 670 / 30 | 670 / 20 | BUV661 |
| | 740 / 35 | 735 / 30 | BUV737 |
| | 820 / 60 | 770 / 40 | BUV805 |
| 405nm | 450 / 50 | 431 / 28 | BV421 |
| | 515 / 20 | 525 / 50 | BV510 |
| | 585 / 15 | 585 / 15 | BV570 |
| | 605 / 40 | 605 / 40 | BV605 |
| | 670 / 30 | 677 / 20 | BV650 |
| | 710 / 40 | 710 / 50 | BV711 |
| | 741 / 40 | 750 / 30 | |
| | 780 / 60 | 770 / 40 | BV786 |
| 488nm | 488/10 | 488 / 10 | |
| | 525 / 50 | 530 / 30 | BB515 |
| | 610 / 20 | 610 / 20 | - |
| | 685 / 35 | 670 / 30 | - |
| | 715 / 30 | 710 / 50 | BB700 |
| | | 750 / 30 | - |
| | 780 / 60 | 770 / 40 | - |
| 581nm | 585 / 15 | 586 / 15 | PE |
| | 610 / 20 | 610 / 20 | PE-Dazzle594 |
| | 670 / 30 | 670 / 30 | - |
| | | 710 / 50 | - |
| | 780 / 60 | 770 / 40 | PECy7 |
| 640nm | 670 / 30 | 670 / 30 | APC |
| | 730 / 35 | 730 / 45 | APC-R700 |
| | 780 / 60 | 770 / 40 | APCFire-750 |

***Supplementary Table 1 Optical configuration of the two cytometers used in this study.*** *Each cytometer was individually QC'ed using different lots of CS&T beads, and 8-peak beads. There was no cross-calibration.*
