## Supplementary Figure 1 for "FlowAtlas.jl: an interactive tool bridging FlowJo with computational tools in Julia"

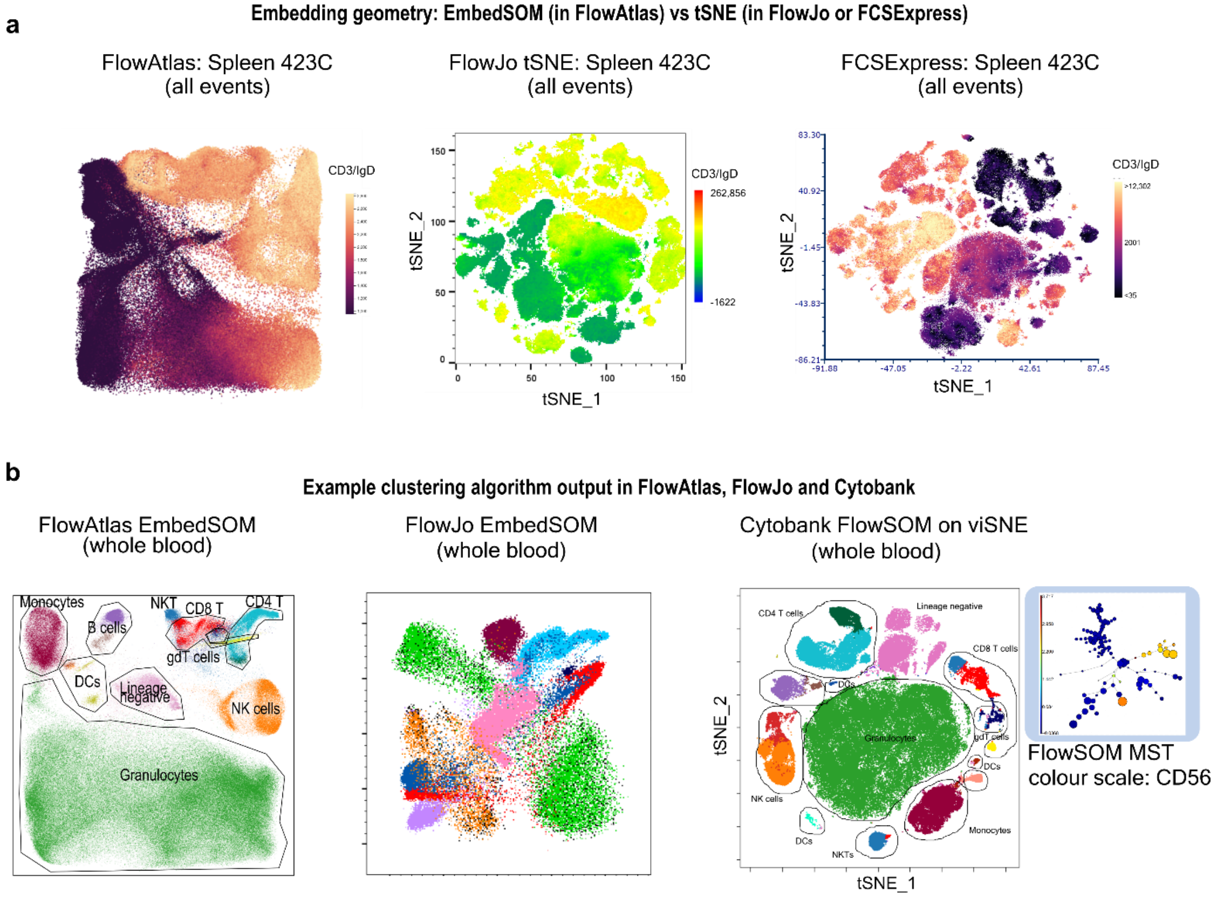


***Supplementary Figure 1 Example graphical output from DR (a) and clustering (b) analysis in different non-command line platforms.*** *(a) DR embedding of a single fcs file (Spleen 423C) from FlowAtlas, FlowJo tSNE and FCSExpress tSNE, coloured by CD3/IgD expression. Embedding scales and heatmap scales are not comparable between plots. (b) Example clustering results from FlowAtlas, FlowJo EmbedSOM plugin and Cytobank applied to a 23-colour spectral cytometry FCS file. Cytobank- inset shows the structure of the resulting minimum spanning tree, MST, which describes relationships between clusters and is coloured by CD56 MFI.*
