## Supplementary Figure 2 for "FlowAtlas.jl: an interactive tool bridging FlowJo with computational tools in Julia"

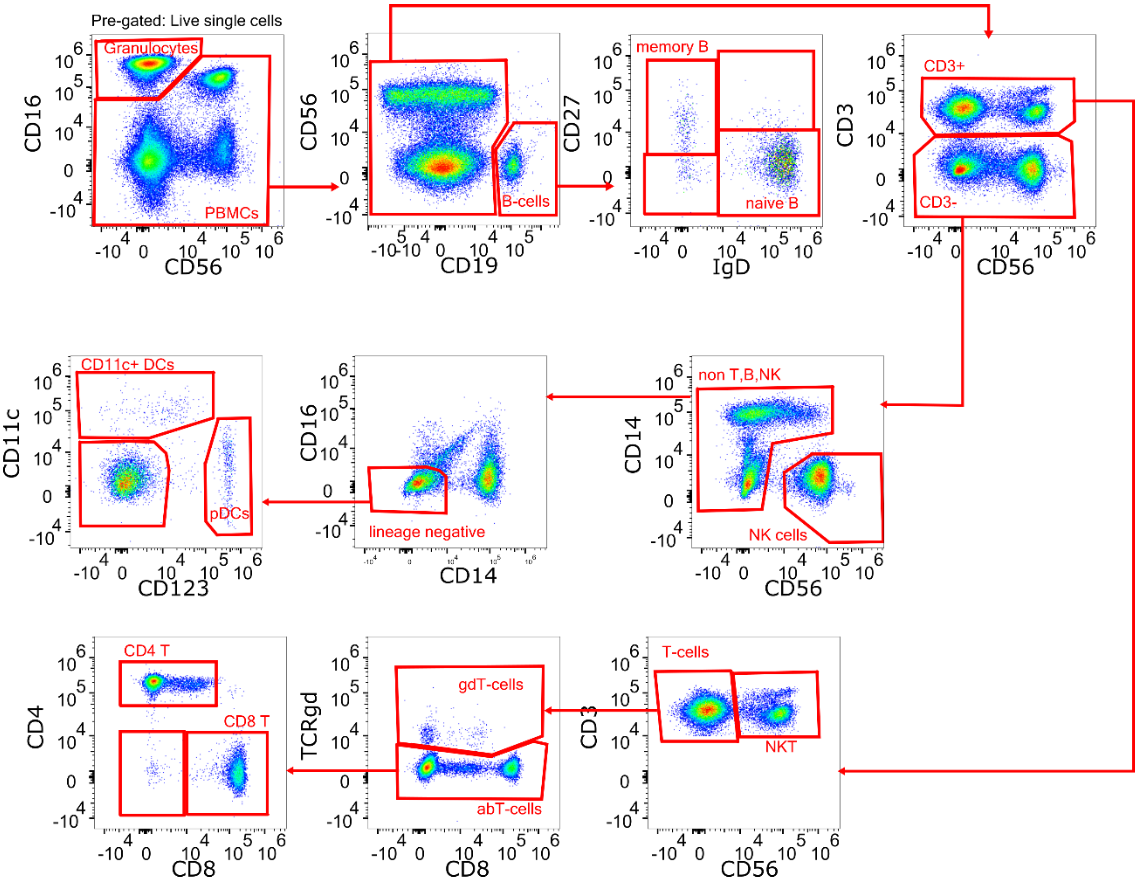


***Supplementary Figure 2 Gating strategy for 23-colour spectral cytometry panel of whole human blood.*** *Debris, dead cells and doublets had already been excluded. Data from Cytobank experiment number 191382 were provided unmixed and compensated. This dataset was used to compare FlowAtlas computational performance at clustering and the workflow for rare population discovery versus Cytobank.*
