## Supplementary Figure 3 for "FlowAtlas.jl: an interactive tool bridging FlowJo with computational tools in Julia"

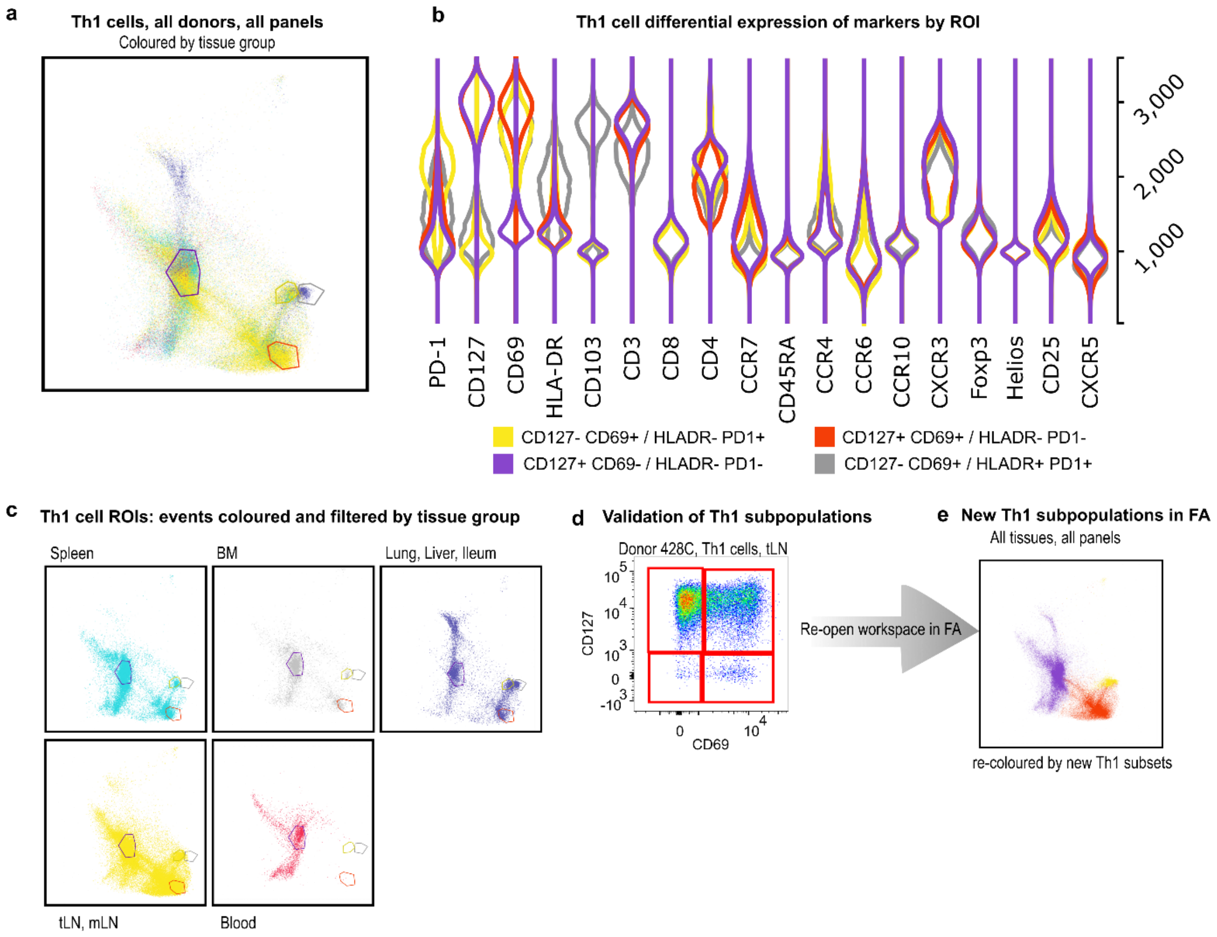


***Supplementary Figure 3 Exploration of Th1 memory cell subsets across tissues and donors in FlowAtlas.*** ***a,*** *Th1 cells were filtered in FlowAtlas, showing all donors coloured by tissue group, and 4 ROIs were drawn. Some tissues were coloured together due to low cell number per sample. (Spleen- cyan; bone marrow- gray; lymph nodes- yellow; non-lymphoid tissues- purple; blood- red)* ***b,*** *Violin plots generated from the 4 ROIs in FlowAtlas. Th1 subsets differed by their expression of CD127, CD69 and PD-1.* ***c****, Th1 cells with the superimposed ROIs, filtered by tissue type, confirmed that CD69+ cells are mainly found in lymphoid and non-lymphoid tissues; the non-lymphoid tissues contained a HLADR+PD1+ population (gray ROI) not present in other tissues. These markers may reflect past cell activation history.* ***d,*** *The new Th1 subsets were validated in FlowJo by creating CD127/CD69 gates in all samples.* ***e,*** *The updated FlowJo workspace was reopened in FlowAtlas to display 3 of the new Th1 suopulations. The HLADR+ PD1+ suopulation is hidden (it is a subset of CD127-CD69+ cells and was not gated in FlowJo; we confirmed it was only present in non-lymphoid tissues).*
