## Supplementary Figure 4 for "FlowAtlas.jl: an interactive tool bridging FlowJo with computational tools in Julia"

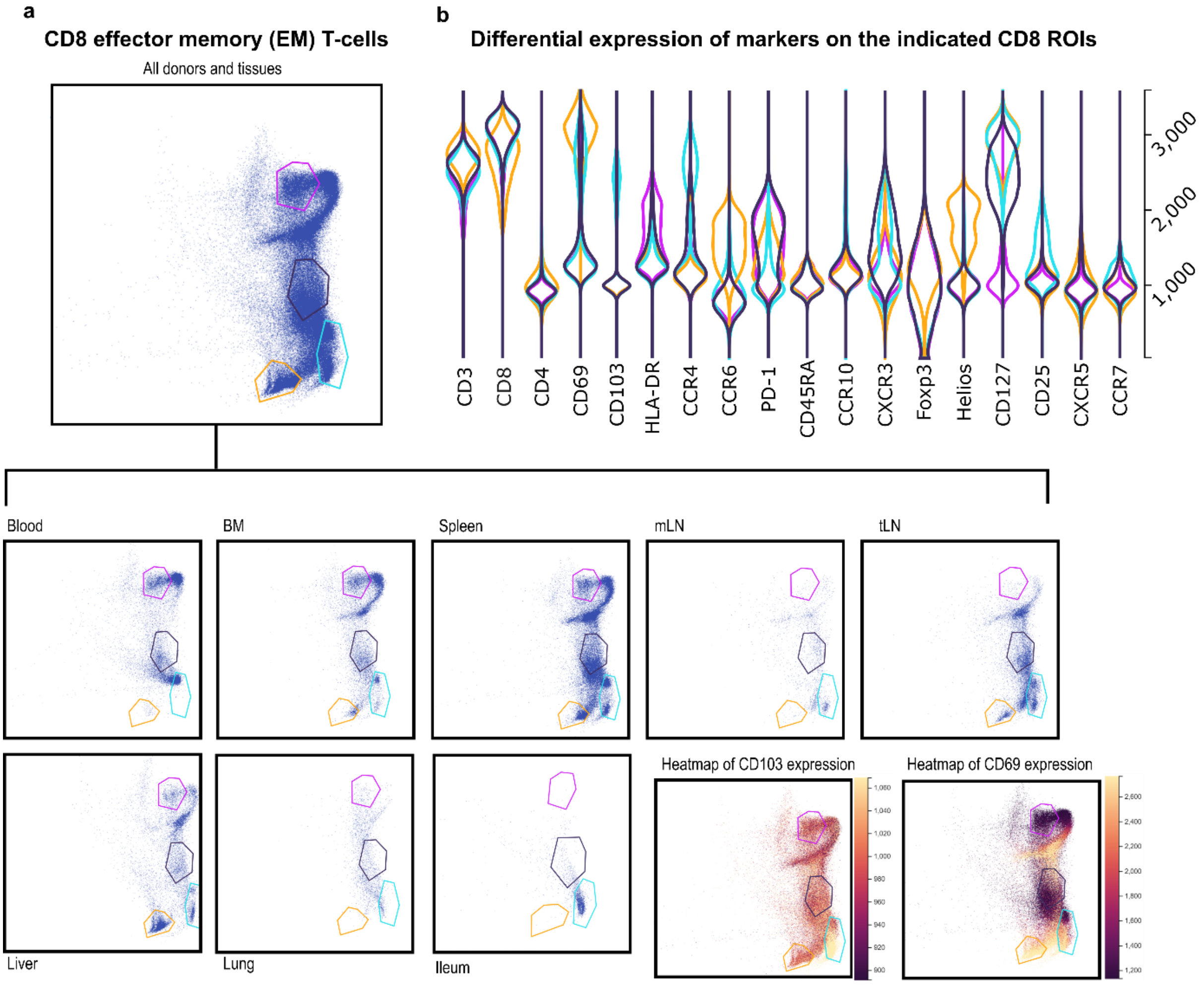


***Supplementary Figure 4 Exploratory analysis of the CD8⁺ effector-memory T cell population.a****,FlowAtlas-generated embedding of CD8⁺ T effector memory (EM) subset, as defined by the initial FlowJo gating strategy, displayed as a composite of all tissues, stained with all panels, and as tissue-specific embeddings. ROIs were drawn around four of many possible subclusters in the composite embedding to auto-generate subcluster-specific violin plots of marker expression. CD69⁺ cells, indicated in cyan and yellow, were largely absent from blood but present in tissues. Although few cells were obtained from the ileum, nearly all co-expressed CD69 and the integrin CD103 as has been described[19],[20]. Heatmaps of CD103 and CD69 expressions are shown on composite embeddings*
