## Supplementary Figure 5 for "FlowAtlas.jl: an interactive tool bridging FlowJo with computational tools in Julia"

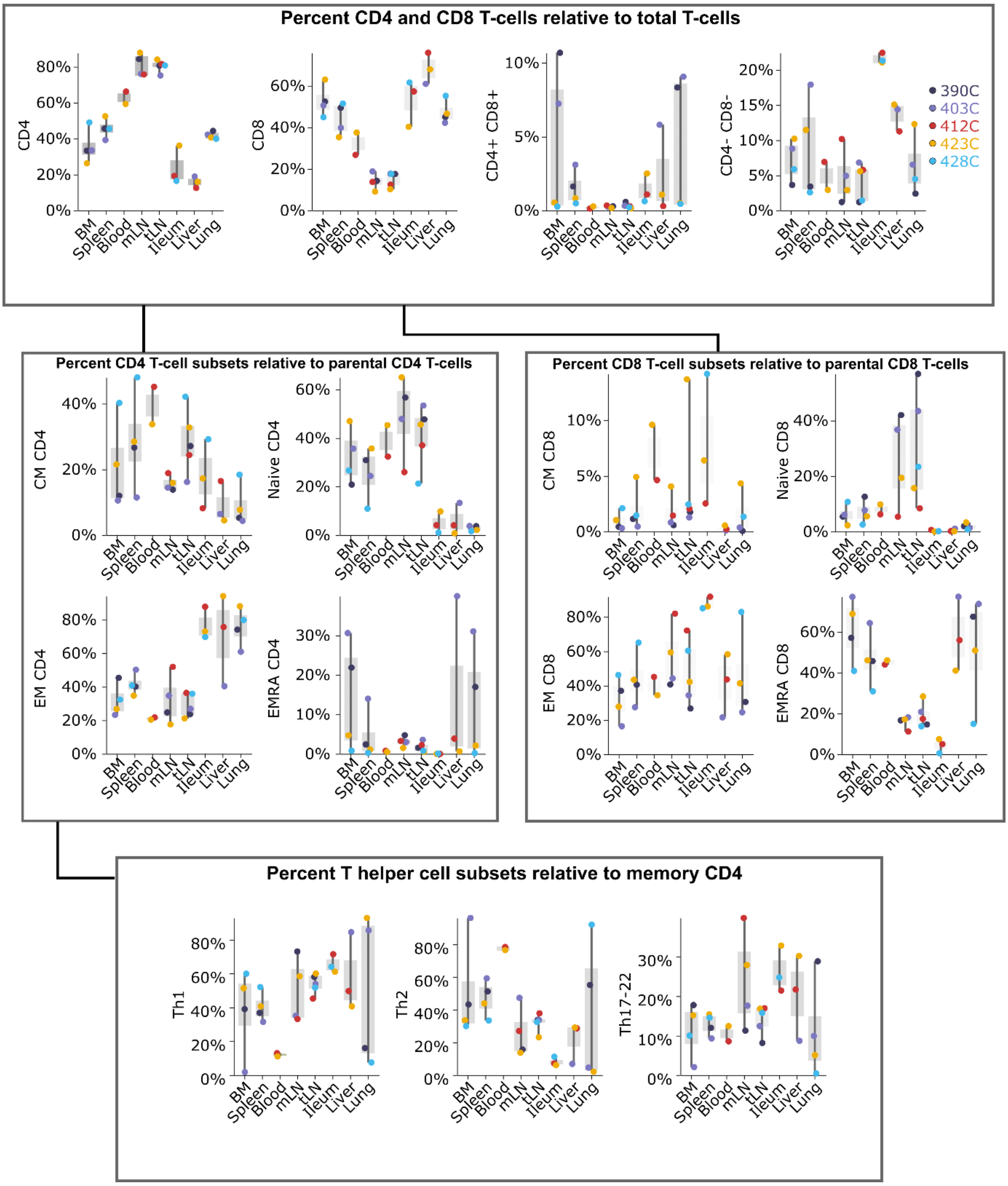


***Supplementary Figure 5 Relative abundance box plots of main T-cell populations in all donors and tissues calculated in FlowAtlas.*** *Blood was available only for donors 412C and 423C. Percent of any combination of populations can be calculated, relative to their sum total. Although not shown, this includes populations which do not share a parent or grandparent gate, such as T-cells and B-cells. CD4 T-cell subsets, especially CM and naïve, were enriched in lymph nodes, whereas ileum, liver, and lung were dominated by CD8 cells, particularly EM and TEMRAs. Within the CD4 memory compartment, frequencies of T helper cell subsets were variable across donors and tissues, with Th2 cells being most frequent in blood. CM - central memory, EM -effector memory; TEMRA - T effector memory cells re-expressing CD45RA*
