## Supplementary Figure 6 for "FlowAtlas.jl: an interactive tool bridging FlowJo with computational tools in Julia"

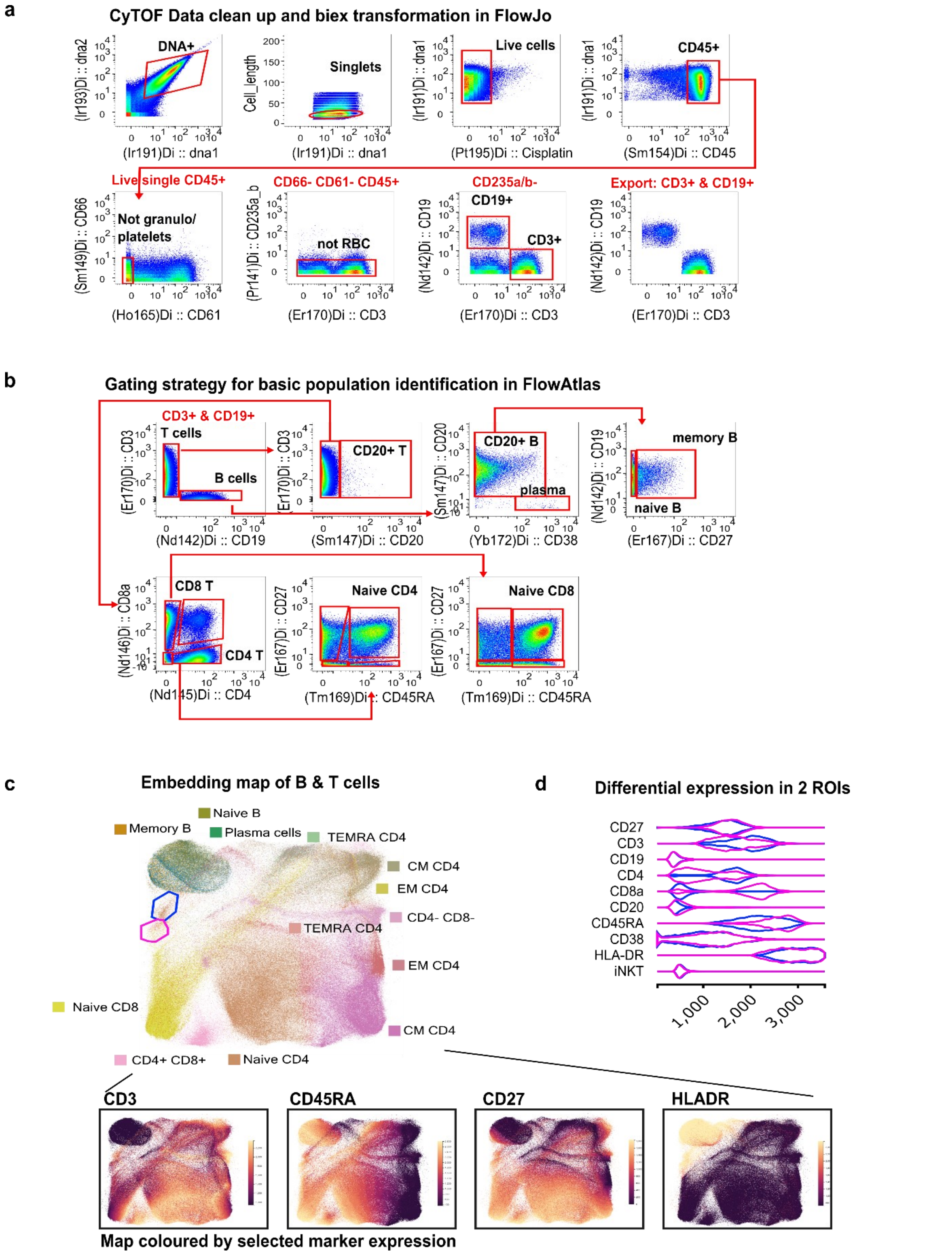


***Supplementary Figure 6 Example analysis of CyTOF data in FlowAtlas. a** Data cleanup to remove irrelevant events, and transformation in FlowJo. **b** Gating strategy for population identification. **c** Embedding in FlowAtlas of exported clean FCS files; inset below- the same embedding coloured by 4 of the markers on the panel. **d** Violin plots of differential expression of markers from two regions in the embedding. Data source: **FlowRepository [FR-FCM-ZZNV](FR-FCM-ZZNV), manuscript** [27933748](27933748) **(Yann Abraham)***
