## Supplementary Figure 7 for "FlowAtlas.jl: an interactive tool bridging FlowJo with computational tools in Julia"

| | Marker | Panel 1 | Panel 2 |
|---|---|---|---|
| | **CD3** | **BUV737** | **BUV737** |
| | **CD4** | BUV395 | Alexa Fluor 700 |
| | CD8 | BV650 | - |
| | CD56 | BV605 | - |
| | CD19 | PECy7 | - |
| | CD16 | Alexa Fluor 700 | - |
| | **CD127** | FITC | PECy7 |
| | **CD45RA** | **BV786** | **BV786** |
| | **CD27** | **V500** | **V500** |
| | **CD25** | PE | BB515 |
| | CD122 | BV421 | - |
| | CR2 (CD21) | - | PECy5 |
| | CD31 | - | BV421 |
| | Ki67 | - | BUV395 |
| | **Foxp3** | Alexa Fluor 647 | PE-Dazzle 594 |
| | Helios | PerCPCy5.5 | |
| Total shared | 7 | 3 | 3 |
| Total unique | 9 | 10 | 7 |
| Total | 16 | 13 | 10 |

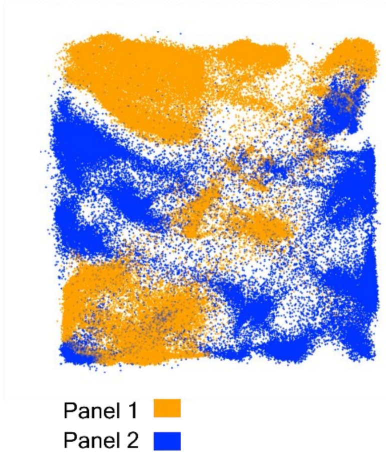


***Supplementary Figure 7 Failure of integration of two highly dissimilar panels due to an insufficient number of overlapping markers and/or fluorochromes.*** *The same donor healthy blood sample was stained during a single experiment with the two indicated panels, using an identical procedure. Data were acquired on the same cytometer with identical settings on the same day. FCS files were cleaned of doublets, dead cells and debris, fully compensated and appropriately transformed prior to embedding. Markers and fluorochromes shared between the panels are indicated in bold.*
